# Key mutations on spike protein altering ACE2 receptor utilization and potentially expanding host range of emerging SARS-CoV-2 variants

**DOI:** 10.1101/2022.04.11.487828

**Authors:** Qiong Wang, Sheng-Bao Ye, Zhi-Jian Zhou, Jin-Yan Li, Ji-Zhou Lv, Bodan Hu, Shuofeng Yuan, Ye Qiu, Xing-Yi Ge

## Abstract

Increasing evidence supports inter-species transmission of SARS-CoV-2 variants from human to domestic or wild animals during the ongoing COVID-19 pandemic, which is posing great challenges to epidemic control. Clarifying the host range of emerging SARS-CoV-2 variants will provide instructive information for the containment of viral spillover. The spike protein (S) of SARS-CoV-2 is the key determinant of receptor utilization, and therefore amino acid mutations on S will probably alter viral host range. Here, in order to evaluate the impact of S mutations, we constructed 20 Hela cell lines stably expressing ACE2 orthologs from different animals, and prepared 27 pseudotyped SARS-CoV-2 carrying different spike mutants, among which 20 bear single mutation and the other 7 were cloned from emerging SARS-CoV-2 variants, including D614G, Alpha (B.1.1.7), Beta (B.1.351), Gamma (P.1), Delta (B.1.135), Lambda (B.1.429) and Mu (B.1.525). Using pseudoviral reporter assay, we identified that the substitutions of T478I and N501Y enabled the pseudovirus to utilize chicken ACE2, indicating potential infectivity to avian species. Furthermore, the S mutants of real SARS-CoV-2 variants comprising N501Y showed significantly acquired abilities to infect cells expressing mouse ACE2, indicating a critical role of N501Y in expanding SARS-CoV-2 host range. In addition, A262S and T478I significantly enhanced the utilization of various mammals ACE2. In summary, our results indicated that T478I and N501Y substitutions were two S mutations important for receptor adaption of SARS-CoV-2, potentially contributing to spillover of the virus to many other animal hosts. Therefore, more attention should be paid to SARS-CoV-2 variants with these two mutations.

## Introduction

COVID-19 epidemic has posed severe threats to global economy and public health, resulting in more than 0.4 billion infections and over 5 million deaths worldwide as reported by World Health Organization (WHO) in 25 February 2022 (https://www.who.int/emergencies/diseases/novel-coronavirus-2019). The disease is caused by severe acute respiratory syndrome coronavirus-2 (SARS-CoV-2), a virus belonging to the genus *Betacoronavirus*, which can infect human and some other animal species, like ferret, dog, tiger and etc [1–3]. During the widespread and long-term transmission in the human population, the virus has been undergoing continuous evolution and adapting to maximize its fitness. Especially, the frequent mutations on the viral spike (S) protein have led to the emergence of viral variants with wider host range, higher transmissibility and elevated immune evasion. First detected in March 2020, the SARS-CoV-2 variants with D614G mutation on S has become the dominant strain globally, tending to almost 100% of prevalence afterwards [4]. Since then, thousands of variants of SARS-CoV-2 have emerged. Currently, WHO has classified several important variants as variants of interest (VOI) and variants of concern (VOC) based on the amino acid changes in the spike protein. VOIs refer to SARS-CoV-2 variants with genetic changes affecting viral transmissibility, disease severity, immune escape, diagnostic or therapeutic escape, and posing apparent epidemiological risk to global public health. VOC are defined as variants not only meeting the definition of VOI but also with detrimental change in COVID-19 epidemiology, virulence or decrease in effectiveness of available diagnostics, vaccines and therapeutics. For instance, B.1.1.7, a typical VOC also known as Alpha, emerged in the United Kingdom in September, 2020, showed high transmissibility and led to a new wave of global epidemic rapidly [5, 6]. Before long, another VOC called B.1.351 (Beta) was detected in South Africa with significantly high resistance to neutralizing antibodies in convalescent sera[7]. P.1 (Gamma), the third VOC, was first detected in Brazil, voiding the plan of herd immunity and overwhelming medical system in Brazil [8]. Later on, VOC Delta (B.1.617.2), firstly detected in India, rapidly spread around the world due to its remarkable transmissibility and ability of immune evasion [9]. The recent VOC B.1.1.529 (Omicron) harboring more than 30 accumulated mutations on its S has spread to more than 80 countries [10]. Besides, two VOI, C.37 (Lambda) and B.1621 (Mu), appeared in Peru and Colombia respectively, fortunately they did not cause any large epidemic [11].

Angiotensin-converting Enzyme 2 (ACE2) is the functional receptor of SARS-CoV-2. Although ACE2 is ubiquitously expressed in a variety of animals, but not all ACE2s can serve as the receptor for SARS-CoV-2 [1, 12]. Our previous study and many other studies showed that the original strain of SARS-CoV-2 was capable of utilizing most mammal ACE2s expect for murine and non-mammal ACE2s [2, 3, 13–18]. However, increasing evidence supports those certain mutations of S expand the host range of SARS-CoV-2, creating new viral reservoirs with the potential to re-infect humans. For example, D614G shifts the S conformation toward an ACE2 binding-competent state, significantly enhancing S-ACE2 affinity, which leads to high infectivity and transmissibility of the virus [19–23]. N501Y, a mutation located in the receptor binding domain (RBD), expands the host range of SARS-CoV-2 to infect mice [24–27] and enhances viral resistance to neutralizing antibodies [28, 29]. Similarly, Q493K and Q498H in RBD significantly increased the binding affinity towards mouse ACE2 [30]. L452R increases viral infectivity and fusogenicity and promotes viral replication [31, 32]. Y453F has been found to increase in hACE2 affinity [33]. N439K and S477N mutations caused immune escape from monoclonal antibodies [34, 35]. P681H mutation in the spike S1/S2 cleavage site, which increase cleavage by furin, potentially impacting the viral cell entry[36]. Nevertheless, most recent studies on S mutations affecting the viral host range are limited to several mammal species, especially on mice. Utilization of non-mammal ACE2 by SARS-CoV-2 variants has been barely investigated. Thus, the host range of the newly emerging SARS-CoV-2 variants may be wider than expected, and underestimation of the viral host range may result in uncontrolled transmission route for the virus.

Assisted by the bioinformatics tool BioAider [37], we continuously monitored the variants of SARS-CoV-2 and screened several hotspot mutations of S, including those carried by VOC and VOI strains, for the study of host adaption. In this study, we used pseudovirus-reporter system to assess 20 ACE2 orthologs of different mammal and avian species for their ability to support viral particle internalization mediated by the SARS-CoV-2 S variants with the hotspot mutations. As the result, we found the impact of S mutations on ACE2 utilization varied among different species. Notably, T478I or N501Y, when individually combined with D614G, enables SARS-CoV-2 to utilize chicken ACE2, indicating that the SARS-CoV-2 variants carrying these mutations might be capable of infecting poultry. In addition, A262S+D614G or T478I+D614G significantly enhanced the utilization of human and monkey ACE2s, potentially promoting viral infectivity to primates. Many other hotspot mutations, in general, increased the utilizing efficiency for ACE2s of domestic animals such as dogs and cats, but reduced that for ACE2s of wild animals, indicating viral adaption to human community during the long-term pandemic. In summary, our study has preliminarily shown the trend of expanding host range and better adaption to human community of SARS-CoV-2, warning a potential concern about more severe prevalence of the virus in the future, though more validation based on living virus is still required to draw a solid conclusion for this issue.

## Results

### Pseudovirus and ACE2 cells

The genomic sequences of human SARS-CoV-2 strains with high coverage scores were downloaded from GISAID database (https://www.gisaid.org/) of which the sampling date was from December 24, 2019 to October 13, 2020. We manually eliminated some strains with poor sequencing quality. A total of 88,247 amino acid (aa) sequences of the S protein were extracted for mutation analyses. Other than D614G which was carried by almost all variants, we screened a total of 20 persistently prevalent and high-frequency single-point mutations, including 11 in the RBD, 7 in the NTD, 1 adjacent to the furin cleavage site (RRAR) and 1 in S2 (Fig 1A and S2 Table). Then, these mutations were introduced to pseudoviral particles carrying SARS-CoV-2 S. In order to validate the successful preparation of these mutant pseudoviruses, the pseudoviral proteins were loaded to SDS-PAGE and subjected to western blot detection of HA-tagged spike proteins. The result showed that all the 20 mutant pseudoviruses were successfully prepared (Fig 1B).

**Fig. 1.**
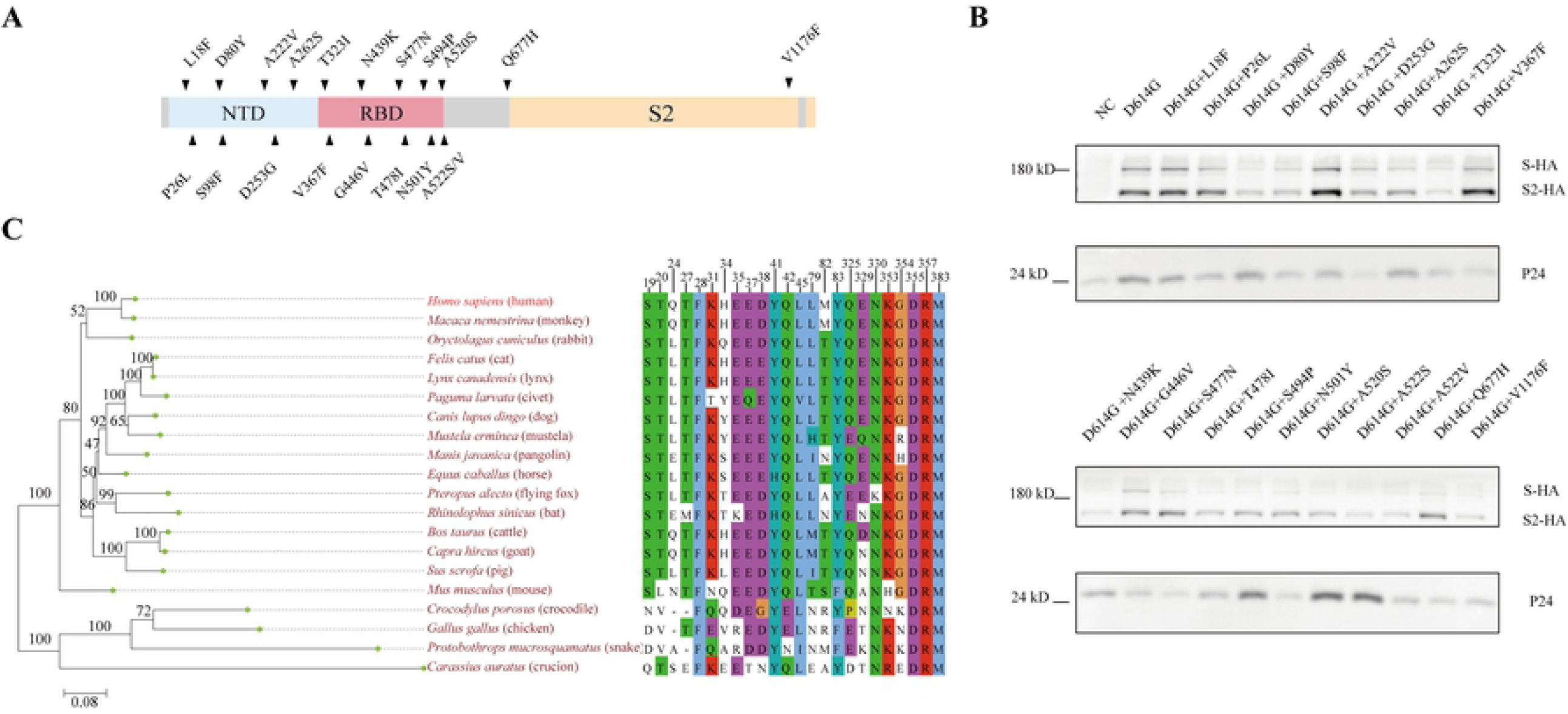
Illustration of 20 single-point spike mutation pseudovirus and Phylogenetic analysis of ACE2 orthologs. (A). Schematic of 20 SARS-CoV-2 spike mutations. (B). Western blot Detection of spike proteins of 20 single-point mutation pseudovirus using an antibody against the HA tag conjugated to the viral spike proteins. HIV-1 p24, a proteins of the carrier pseudovirus, was detected as the loading control. **C**. The sequences of 20 ACE2 proteins were analyzed and the phylogenetic tree was built. The residues of human ACE2 at the SARS-CoV-2 RBD/ACE2 interface are shown.

Phylogenetic analysis was performed for the 20 ACE2s of different animal species and showed that they could be branched into two major groups. The first group consists of mammals which could be further divided into primate and non-primate; the second group includes avian, reptile and fish (Fig 1C, left panel). Additionally, we aligned the 24 key aa residues of ACE2 located at the domain interfacing SARS-CoV-2 S protein (Fig 1C, right panel). Of note, most mammalian ACE2s were highly conserved on these aa residues with human ACE2, except mouse ACE2 which differed in aa residues located at position 31, 83 and 353. In comparsion, these residues in non-mammalian ACE2s were less conserved.

### T478I and N501Y enabled SARS-CoV-2 S to utilize chicken ACE2s

Our previous study showed that the original SARS-CoV-2 S could not utilize avian ACE2, but SARS-CoV can [13]. Here, we tested the ability of the 20 ACE2 mutants to utilize chicken ACE2. Cells expressing chicken ACE2 showed a strong luminescence signal when infected by the pseudoviruses carrying T478I+D614G or N501Y+D614G compared to the control carrying D614G only, indicating that the substitutions of T478I and N501Y enabled the utilization of chicken ACE2 (Fig 2A). Interestingly, the 478th aa residue in RBD of SARS-CoV is K, which is consistent with Delta and Omicron variant (Fig 2B).

**Fig. 2.**
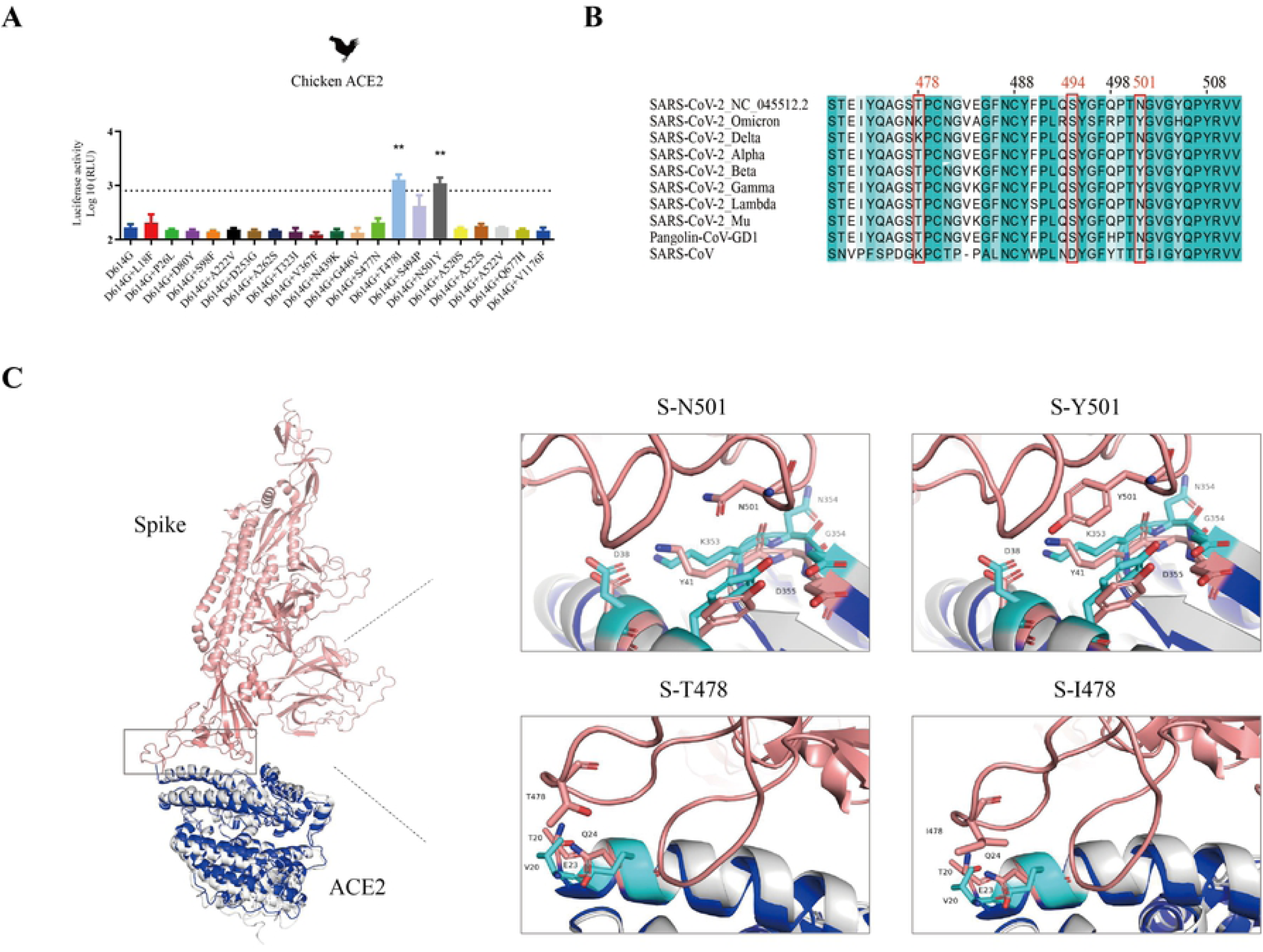
The characteristic of SARS-CoV-2 T478I and N501Y mutations. (A) Hela cells expressing chicken ACE2 were infected by 20 SARS-CoV-2 spike mutations pseudoviruses. At 48 h post infection, pseudovirus entry efficiency was determined by measuring luciferase activity in cell lysates. The results were presented as the mean relative luminescence units (RLUs) and the error bars indicated the logarithm (base 10) of the standard deviations of the RLUs (n= 9). (B) Sequence alignment of the RBD sequences from the SARS-CoV-2, SARS-CoV-2 variants and SARS-CoV. (C) Structural models of N501Y and T478I were generated based on the crystal structure of the SARS-CoV-2 S/ACE2 complex. SARS-CoV-2 S, chicken ACE2 and human ACE2 were colored in pink, gray and blue, respectively. Residues involved in the interaction are labeled.

To test whether position 478 and 501 of S influenced chicken ACE2 usage efficiency, we used homology-based structural modeling to analyze the effects of residue substitutions at position 478 and 501. Structural models of T478I and N501Y were generated based on the crystal structure of the SARS-CoV-2 S/ACE2 complex. We found that position 478 and 501 were located at the interaction domain with chicken ACE2. Furthermore, in contact with 501 of S, the interface between chicken ACE2 and human ACE2 consists 5 residues (D38, Y41, K353, N354, D355 in chicken ACE2, D38, Y41, K353, G354, D355 in human ACE2). When 501 is N, N354 of chicken ACE2 disrupts N501 forming a hydrogen bond with K353. When 501 is converted to Y, it is closer to ACE2 353 and 41, similar to human ACE2, forming an additional π-π interaction with Y41 and an additional π-cation interaction with K353 (Fig 2C). In contact with 478 of S, the interface between chicken ACE2 and human ACE2 consists 2 residues (V20, E23 in chicken ACE2, T20, Q24 in human ACE2). In short, T478I and N501Y on S mutations could expand host range to birds.

### Hotspot mutants of SARS-CoV-2 S tended to utilize the ACE2 of primates and domestic animals

In the test of mammalian ACE2, we found that the T478I or A262S significantly increased ability of pseudovirus to utilize human and monkey ACE2 compared to D614G only control, indicating these two mutants enhance the utilization of primate ACE2s (Fig 3A and 3B). However, P26L, A522S/V and Q677H significantly reduced the utilization of human ACE2, but showed no change for monkey ACE2 (Fig 3A and 3B).

**Fig. 3.**
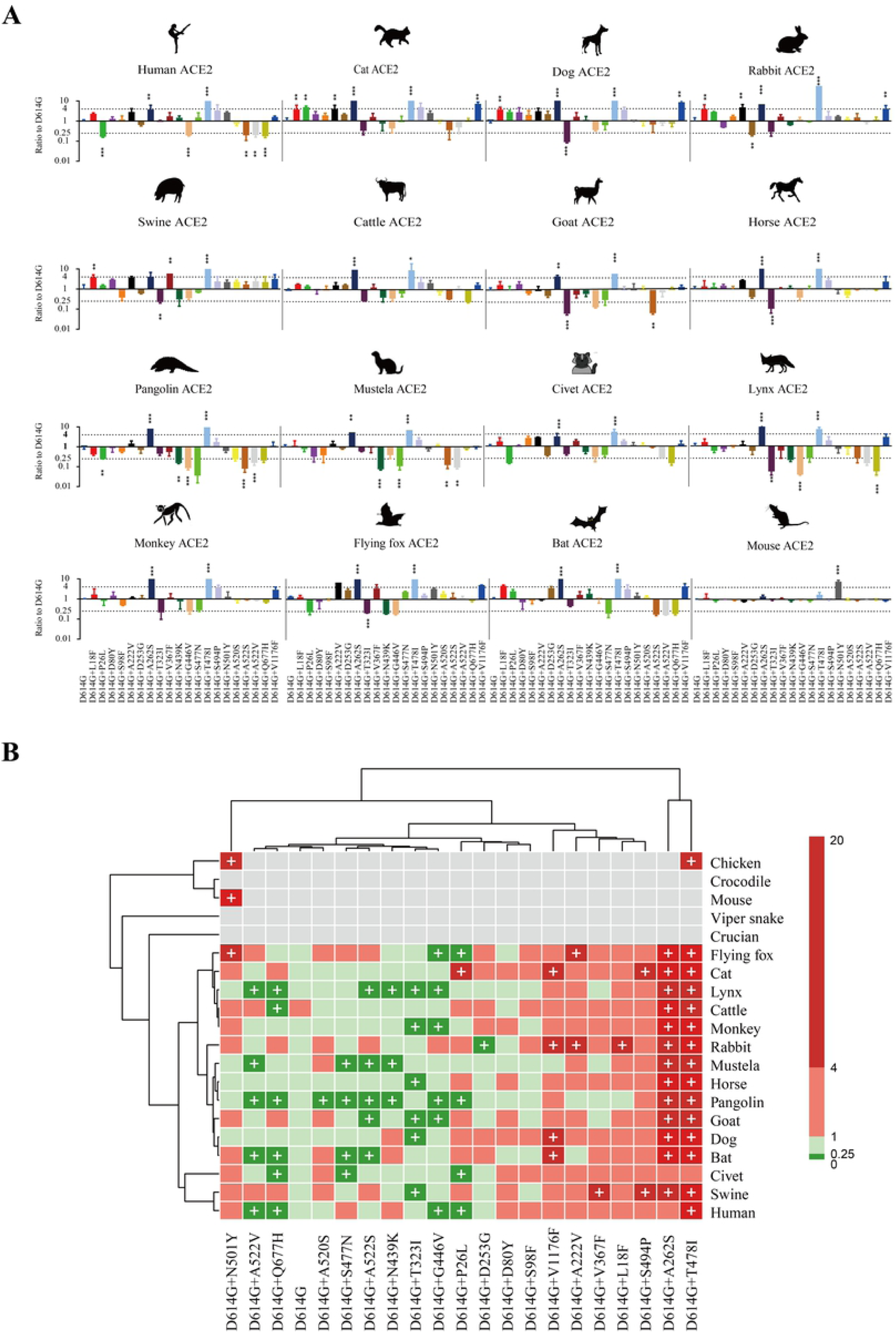
Infectivity of SARS-CoV-2 mutations in cell lines expressing ACE2 proteins from 20 different species. (A) Equal amounts of overexpression plasmids carrying ACE2 orthologs from different species were transfected into Hela cells. Then, Cells were infected with the same copy number of SARS-CoV-2 variants after quantification. luminescence signals were collected 48 h later. Ratios between mutants and the D614G reference strain were calculated. The results were obtained from three independent experiments and the values shown indicate means ± SEMs. Dashed lines indicate the threshold of fourfold difference. Asterisks indicate statistical significance. (B) The heatmap was created by pheatmap package in R program. The color of the square indicates different ratio ranges value, gray indicates that it cannot be infected. red indicates increased infection compared to D614G, green indicates reduced infection compared to D614G, plus represents the threshold of fourfold difference. The row clustering and column clustering through “complete” methods in pheatmap package, and the distance type was set to “correlation”.

For non-primate mammals, our test showed that L18F, A222V, A262S, T478I and V1176F increased usage efficiency of dog and cat ACE2s more than 4 folds (Fig 3A and 3B). Since dogs and cats are pet animals, we guessed that the hotspot mutations might enhance the utilization of ACE2s of domestic animals generally. This speculation was verified in the following tests. Swine, cattle, goat and horse are common domestic animals farmed in human communities, and both A262S and T478I mutations significantly increase the utilization of ACE2s of these farm animals (Fig 3A and 3B). Especially, even more mutations, including L18F, A222V and V367F, significantly increased the utilization of swine ACE2. In summary, these results implied a potential of SARS-CoV-2 adaption to domestic animals caused by the hotspot mutations of S.

In comparison, the above mutations, except A262S and T478I, posed little influence on the utilization of ACE2s of wild animals. On the contrary, most mutants, including N439K, G446V, S477N, A522V and Q677H, significantly reduced ACE2 utilization efficiency for various wild animals such as pangolin, mustela and lynx, suggesting that these animals may be one-time intermediate hosts, not adapted to the evolution of SARS-CoV-2. Furthermore, P26L, T323I, N439K and G446V led to more than 4-fold reduction in the utilizing efficiency of flying fox ACE2, while S477N, A522S/V and Q677H decreased the utilization of bat ACE2 in bats. Although a few mutations, such as A262S and T478I, enhanced the utilization of wild animal ACE2s, the number of such mutation was much fewer than those for domestic animals. Therefore, the SARS-CoV-2 mutants seemed to lower their adaption to wild animals.

### Host range shift of VOC and VOI variants

Next, we evaluated the potential host range shift of current VOC and VOI variants by the pseudovirus reporter assay. Our results showed that the S with D614G and of Lambda variants significantly enhanced ability to utilize mammalian ACE2, compared to the original S. Beta and Mu variants significantly increased the utilization of ACE2 of monkeys, indicating that these variants may have enhanced infectivity in primate animals. In addition, Mu variant has an increased ability to utilize flying fox ACE2, which is consistent with N501Y point-mutant. Besides, the Alpha, Beta, Gamma and Mu variants, comprising N501Y mutation, display significantly acquired abilities to infect mouse ACE2-overexpressing cells (Fig 4A, B). This suggested that SARS-CoV-2 variants containing the N501Y mutation could acquire the ability to transmit human to mouse and can be transmitted among mouse. Whereas, variants with the T478K or N501Y mutation unable to utilize chicken ACE2 for cell entry. These variants were also unable to utilize fish and reptilian ACE2s, as did the 20 mutants above (S1 and S2 Figs).

**Fig. 4.**
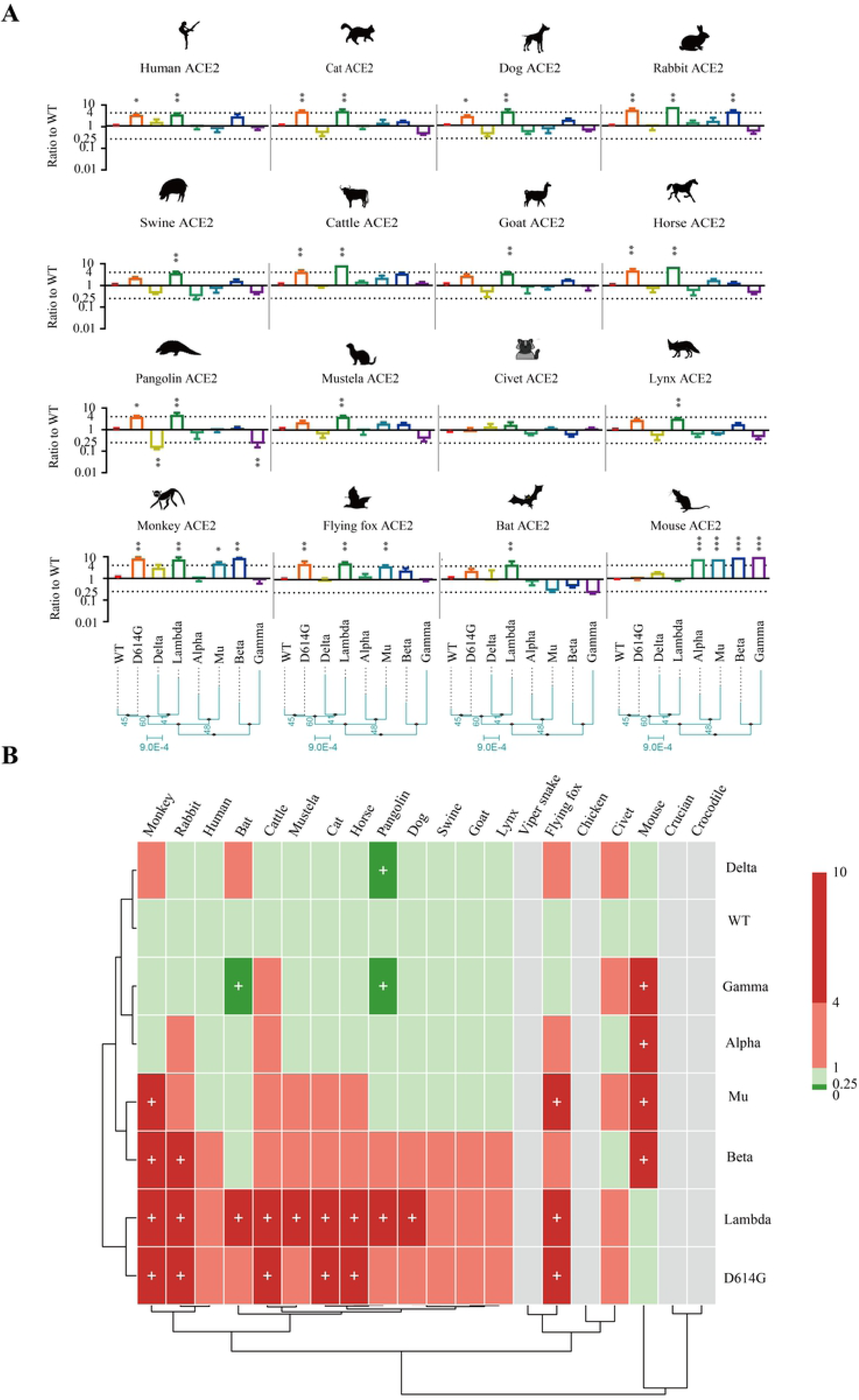
Animal tropism of VOC and VOI variants. (A) Equal amounts of overexpression plasmids carrying ACE2 orthologs from different species were transfected into Hela cells. Then, Cells were infected with the same copy number of SARS-CoV-2 variants after quantification. luminescence signals were collected 48 h later. Ratios between mutants and the WT reference strain were calculated. The results were obtained from three independent experiments and the values shown indicate means ± SEMs. Dashed lines indicate the threshold of fourfold difference. Asterisks indicate statistical significance. (B) The heatmap was created by pheatmap package in R program. The color of the square indicates different ratio ranges value, gray indicates that it cannot be infected. red indicates increased infection compared to WT, green indicates reduced infection compared to WT, plus represents the threshold of fourfold difference. The row clustering and column clustering through “complete” methods in pheatmap package, and the distance type was set to “correlation”.

## Discussion

The persistent emergence of diverse SARS-CoV-2 variants which are mainly defined by S mutations has been posing great challenges to the control of the current COVID-19 pandemic. Viral spillover from animals to humans and further spillback to other animals were believed to be a key factor for the frequent emergence of SARS-CoV-2 variants [14, 33]. The host range of SARS-CoV-2 is mainly determined by the recognition between S protein and its cellular receptor ACE2 orthologs. Genetic changes in the S usually switch viral host range and alter viral infectivity [26, 38]. Predicting and monitoring the interaction among different S mutants and ACE2 orthologs are critical to identified animal reservoirs and intermediate hosts of SARS-CoV-2, which will help to prevent the spread and circulation of SARS-CoV-2.

Birds are important reservoirs of a variety of viruses, including some coronavirus and influenza virus[39]. However, the original strain of SARS-CoV-2 was unable to utilize avian ACE2 and replicate poorly in poultry [2]. In our study, we assessed the interaction of 27 S mutants to 20 ACE2 orthologs using pseudovirus. Interestingly, T478I and N501Y mutations on S enabled the pseudovirus to utilize chicken ACE2, indicating potential expansion of SARS-CoV-2 host range to avian species. This result accorded with the latest report that Omicron variant which carrying N501Y mutation was capable of infecting cells expressing avian ACE2 [10]. Notably, N501Y mutation is widely exist in other VOC and VOI variants, such as Alpha, Beta, Gamma and Lambda [6, 36]. T478I showed an even higher frequency in our dataset which was carried by more than two million variants and about 2-fold of N501Y, and the VOC of Delta and Omicron carry T478K which may be similar to T478I in altering ACE2 utilization (S2 Table). These results suggested that 478^th^ and 501^th^ could be potentially key site for interspecies transmission of SARS-CoV-2 and raised the concern about the neglecting of circulation of SARS-CoV-2 in poultry and wild birds.

The change of ACE2 utilization caused by N501Y and T478I mutations may be explained from the view of protein structure. Previous analysis of the S-human-ACE2 complex revealed that N501Y probably introduced additional π-π and π-cation interactions between S RBD and an interface of human ACE2, increasing the binding affinity [40, 41]. According to the structure stimulation of S-chicken-ACE2 complex, this interface showed a similar structure to that of human ACE2, providing the structural basis for cross-species interaction of SARS-CoV-2 variants. However, in chicken ACE2, position 354 is an Asn not a Gly, and D354 in chicken ACE2 does not fit into RBD as tightly as G354 of hACE2. Therefore, more verification is required to confirm the utilization of chicken ACE2 by N501Y S. T478 is located at the RBD-ACE2 binding interface close to the N-terminal T20 of human ACE2[42]. T478I substitution brought a slightly larger side chain and changed S protein conformation, which might allow interaction to V20 of chicken ACE2 and enhanced the stabilization of the RBD-ACE2 complex. Considering chicken is an important agricultural animal widely kept in human communities, the detection of SARS-CoV-2 in farmed chickens may be needed for epidemic control [43].

Old world monkeys were used as experimental animal models for SARS-CoV-2 infection due to their similarity to human in ACE2 sequences [44]. However, our study demonstrated that P26L, A522S/V and Q677H significantly reduced the utilization of human ACE2, but showed no change for monkey ACE2 (*Macaca nemestrina*), regardless of the high identity of monkey ACE2 and human ACE2. This indicated that sequence identity did not necessarily correlate the ability of efficiency of receptor utilization by SARS-CoV-2. This speculation is supported by a previous study showing that three new world monkey species did not support SARS-CoV-2 entry due to two specific residues (H41 and E42) within their ACE2[12].

Domestic animals (pets and livestock) frequently contact with humans which may be intermediate hosts of SARS-CoV-2. Several studies demonstrated that SARS-CoV-2 has been detected in cats and dogs, and laboratory experiments revealed that SARS-CoV-2 replicated efficiently in cats[45–48]. Our results showed that L18F, A222V, A262S, T478I and V1176F increased S utilization of dog and cat ACE2s. Thus, the variants carrying these mutations may spread among pet animals, which should be paid special attention in the epidemic control.

Natural SARS-CoV-2 infections in wild animals have been reported in tiger, lions, and white-tailed deer [16, 49]. Especially, pangolins carrying the SARS-CoV-2-like CoVs manifested clinical symptoms and histological pathogenicity [50]. However, most S mutations, including N439K, G446V, S477N, A522V and Q677H, significantly reduced ACE2 utilization efficiency for various wild animals such as pangolin, mustela and lynx. Similar results were also found in our study for bat ACE2s. Bats are highly diverse with the second largest number of species in mammals, carrying a wide variety of coronaviruses [51]. Bats might be potential hosts of ancestral SARS-CoV and SARS-CoV-2[1, 52]. Several studies showed that *Rhinolophus sinicus* could support SARS-CoV-2 entry, whereas its congeneric relatives *Rhinolophus ferrumequinum* and *Rhinolophus pearsonii* could not [53].As shown in our study, ACE2 mutants containing P26L, T323I, N439K and G446V dramatically decreased the utilization of flying fox (*Pteropus Alecto*) ACE2, but had no obvious effects on bat (*Rhinolophus sinicus*) ACE2. Furthermore, S477N, A522S/V and Q677H decreased the utilization of bat ACE2, whereas had no obvious effects in flying fox ACE2. Overall, these results suggest that the viral has altered its adaptation to human communities during the epidemic and wild animals can hardly serve as intermediate hosts during the viral spread in human communities.

Recent studies reported that Omicron variants infected mice, accumulated mutations when circulating in mice, and then re-infected humans [54]. Therefore, rodents may be important intermediate hosts for the transmission of SARS-CoV-2 currently [18]. According our results, N501Y S mutant could utilize mouse ACE2, consistent with previous reports [24, 55, 56]. L18F, A222V, A262S, T478I and V1176F could promote the utilization of rabbit ACE2 (Fig 3). Notably, these mutations are widely carried by VOC and VOI variants. For instance, L18F, P26L and V1176F mutations are carried by Gamma variant; D80Y is present in Beta and Delta; and T478K is carried by Omicron variant; N501Y mutation widely exists in current pandemic variants N501Y mutation is widely exist in VOC and VOI variants, such as Alpha, Beta, Gamma, Omicron and Lambda (S3 Fig). Therefore, in addition to poultry and pets, rodents in the communities may also be important intermediate for the spreading and reservoirs for the mutations of SARS-CoV-2

In summary, we provided a comprehensive view of potential host range change of SARS-CoV-2 caused by S mutations using pseudovirus reporter assay. Notably, T478I and N501Y mutations conferred the ability to utilize chicken ACE2, probably extending the host range of SARS-CoV-2 to birds. Most mutations favored the utilization of domestic mammal ACE2s but weakened that of wild animal ACE2s, indicating the trend of SARS-CoV-2 to adapt human community during the long-term transmission among humans. The shift of viral host range may provide more routes for viral transmission and more reservoirs for viral evolution against vaccines or drugs, which may explain the frequent re-emergence of COVID-19 epidemic around the world even when most population has been vaccinated. Thus, monitoring the host range shift of newly emerging SARS-CoV-2 variants is critical for epidemic control. Nevertheless, our results were mainly based on the pseudovirus assay which might contain potential bias, and thus further verification based on living SARS-CoV-2 variants is required to confirm these findings.

## Methods and materials

### Cell lines and plasmid construction

HEK293T and Hela cells were maintained in Dulbecco’s modified Eagle’s medium (DMEM) supplemented with 10% fetal bovine serum (FBS), 100 units of penicillin and 0.1 mg/ml of streptomycin in 5% CO2 at 37 °C. Full-length SARS-CoV-2 spike (GenBank accession number: MN908947.3), Variants Alpha, Beta, Gamma, Delta, Lambda and Mu spike (GISAID accession number: EPI_ISL_5253387, EPI_ISL_5195381, EPI_ISL_5254522, EPI_ISL_3558827, EPI_ISL_4348182, EPI_ISL_431119) were all synthesized and subcloned into the pcDNA3.1 vectors with a C-terminal HA tag.

Spike point-mutated plasmid constructed by site-directed mutagenesis. pcDNA3.1-S2 plasmid was used as the template to generate the plasmid with mutagenesis in S gene. Following procedure of circular PCR, 15 to 20 nucleotides before and after the target mutation site were selected as forward primers, while the reverse complementary sequences were selected as reverse primers. Following PCR, the template chain was digested using DpnI restriction endonuclease (NEB, USA). Afterward, the PCR digested product was directly used to transform E. coli Top 10 competent cells, single clones were selected and then sequenced. The primers designed for the specific mutation sites are listed in S1 Table

### Western blot analysis

The spike protein of SARS-CoV-2 or variants on pseudovirions were detected by using western blot. Briefly, to pellet down pseudovirions, the viral supernatants were centrifuged at 30,000 rpm for 2 h at 4 °C through a 20% sucrose cushion, and virus pellets were resuspended into 100 μL PBS. The samples were boiled for 10 min and separated in a 10% SDS-PAGE gel and transferred to 0.45 μm nitrocellulose membrane (Millipore, USA). For detection of S protein, the membrane was incubated with anti-HA tag mouse monoclonal antibody (bimake, USA,1:2000), and the bound antibodies were detected by Horseradish Peroxidase (HRP)-conjugated goat anti-mouse IgG (Abbkine, China, 1:5,000). For detection of HIV-1 p24 in supernatants, monoclonal antibody against HIV p24 (p24 MAb) was used as the primary antibody at a dilution of 1:8,000, followed by incubation with HRP-conjugated goat anti-mouse IgG at the same dilution.

### Preparation of pseudovirus

The preparation of pseudovirus was performed based on our previous study. In brief, one day prior to transfection for virus production, HEK293T cells were digested and adjusted to an amount of 7×10^6^ cells in a 10cm culture medium and incubated overnight in an incubator at 37 °C with 5% CO2. When cells reached 80%-90% confluence, HEK293T cells were co-transfected with a luciferase-expressing HIV-1 plasmid (pNL4-3.Luc.R-E-) and a plasmid encoding HA-tagged SARS-CoV-2 spike, SARS-CoV-2 variants spike or point-mutated spike, according to the instructions of Lipofectamine 2000 (Invitrogen). After 4-6 hours, the cell medium was discarded, and 10 mL fresh complete DMEM was added to the 10cm cell dish. After 48h or 72h, the supernatant containing pseudoviruses were harvested, filtered, aliquoted, and frozen at −80°C for future use.

### Pseudovirus quantification and infection assay

Before infection, all the types of pseudovirus were quantificated by RT-PCR. Generally, RNA of SARS-CoV-2 S, SARS-CoV-2 viriants S and point-mutated pseudovirus were extracted using the QIAamp Viral RNA Mini Kit (QIAGEN, Cat#52906), and served as template for reverse transcription using the TransScript All-in-One First-Strand cDNA Synthesis SuperMix for qPCR reagent (TransGen Biotech, Cat#AT341-02). Virus quantification by real-time PCR was performed using the QPCR Mix (Bimake, Cat#B21202), following the supplier’s instruction. The P24 gene of HIV virus was cloned into the vector pCDNA3.1(+) as a plasmid standard, with the viral copy number calculated accordingly.

In infection assay, after quantitative analysis by RT-PCR, the pseudovirus were diluted to the same particle number, and 100μL was taken and added into a 96-well cell culture plate. A total of 20 different cell lines, including different ACE2-overexpressing cells were digested by trypsin and 2×10^4^ cells were added to each well of the 96-well plate. Then, after incubation in a 37 °C for 48 h. Cells were then washed with PBS buffer and lysed. Lysates were tested for luciferase activity (Promega, USA). Each infection experiment was carried out on for three times.

### Phylogenetic analysis and sequence alignment

The ACE2 aa sequences were aligned by MAFFT v7.149 in BioAider. Then we constructed the maximum likelihood phylogenetic tree of ACE2 by IQ-tree v1.6.10 program with 10,000 ultrafast bootstraps (https://academic.oup.com/mbe/article/32/1/268/2925592), and the most appropriate evolutionary model was JTTDCMut+G4 which calculated using ModelFinder according to the bayesian information criterions.

### Structural models

The hACE2/S complex was used as a template for homology modeling[42]. The mutations in the models were aligned, and the interactions between the SARS-CoV-2 S and ACE2 proteins were compared in PyMOL.

## Acknowledgements

We thank all study participants for their generous participation and contribution to this work.

## Author contributions

**Conceptualization:** Qiong Wang, Ye Qiu, Xing-Yi Ge.

**Data curation:** Qiong Wang, Sheng-bao Ye.

**Formal analysis:** Qiong Wang, Ye Qiu.

**Funding acquisition:** Xing-Yi Ge, Shuofeng Yuan.

**Supervision:** Xing-Yi Ge.

**Validation:** Qiong Wang, Sheng-bao Ye.

**Visualization:** Qiong Wang, Zhi-Jian Zhou.

**Writing-original draft:** Qiong Wang

**Writing-review & editing:** Qiong Wang, Jin-Yan Li, Ji-Zhou Lv, Bodan Hu, Shuofeng Yuan, Ye Qiu, Xing-Yi Ge.

## Supporting information

**S1 Fig. Entry efficiency of different S mutants pseudoviruses into ACE2-expressing cells**. HeLa cells expressing different ACE2 orthologs were infected by SARS-CoV-2 variants pseudoviruses. At 48h post-infection, pseudovirus entry efficiency was determined by measuring luciferase activity in cell lysates. The results were presented as the mean relative luminescence units (RLUs) and the error bars indicated the logarithm (base 10) of the standard deviations of the RLUs (n= 9).

**S2 Fig. Entry efficiency of SARS-CoV-BJ01, SARS-CoV-2 variants pseudoviruses into ACE2-expressing cells**. HeLa cells expressing different ACE2 orthologs were infected by SARS-CoV-2 variants pseudoviruses. At 48h post-infection, pseudovirus entry efficiency was determined by measuring luciferase activity in cell lysates. The results were presented as the mean relative luminescence units (RLUs) and the error bars indicated the logarithm (base 10) of the standard deviations of the RLUs (n= 9).

**S3 Fig. Schematic of SARS-CoV-2 variants**. The most representative amino acid mutations of each variant were marked in this study. Each mutation in each SARS-CoV-2 variant is indicated relative to the reference WT sequence. NTD N-terminal domain, RBD Receptor-binding domain, RBM Receptor-binding motif, HR1 Heptad repeats 1, HR Heptad repeats 2.

**S1 Table. Primers design for mutagenesis**.

**S2 Table The information of SARS-CoV-2 mutants**.

